# T cell receptor sharing in hypersensitivity pneumonitis

**DOI:** 10.1101/2024.11.13.623392

**Authors:** Wezi Sendama, Wendy Funston, Richard CH Davidson, Anthony J Rostron, A John Simpson

**Author notes:** **Corresponding author:** Wezi Sendama, 2^nd^ floor William Leech Building, Medical School, Framlington Place, Newcastle upon Tyne NE2 4HH. These authors contributed equally (Anthony J Rostron, A John Simpson). **Author contributions:** WS devised the study and performed analysis; WS, WF, RCHD, AJR and AJS interpreted results and drafted the manuscript. **Funding support:** WS is a National Institute for Health and Care Research (NIHR) Academic Clinical Lecturer. AJS is an NIHR Senior Investigator. The views expressed are those of the authors and not necessarily those of the NIHR or the Department of Health and Social Care.

## Abstract

Hypersensitivity pneumonitis (HP) is characterised by an excessive pulmonary T cell response in susceptible individuals after exposure to inhaled antigens. The most effective treatment for the condition is antigen avoidance, but in most cases an antigen cannot be identified. Profiling antigen-specific T cell responses may form the basis of a strategy to identify causative antigens of the disease. We used public RNA sequencing data and reconstructed T cell receptor (TCR) repertoires from blood and bronchoalveolar lavage samples from patients with HP, idiopathic pulmonary fibrosis and healthy controls. After excluding TCR sequences likely to be related to common microbial exposures in patients with HP, we identified TCRs shared between patients with shared human leukocyte antigen alleles, indicating patients likely to have a common causative antigen. We also identified clusters of TCRs containing identical and similar TCR clones in individual patients that are plausibly related to the causative antigens in those patients. These results establish the feasibility of profiling TCR repertoires to identify antigens in HP.

## Introduction

Hypersensitivity pneumonitis (HP) results from a disordered pulmonary immune response to antigens inhaled by a susceptible individual. The antigens that can be associated with the development of HP are diverse.^1^ A key pillar of disease treatment is antigen avoidance, but it is also the case that in up to 60% of cases a causative antigen cannot be identified despite rigorous efforts.^2^ Failure to identify a causative antigen is associated with poorer disease outcomes.^3^

The pulmonary immune response in HP is characterised by a lymphocyte-rich alveolitis, and the cellular portion of bronchoalveolar lavage (BAL) fluid from patients with HP can comprise up to 80% lymphocytes (compared to 15% or lower in control subjects).^1^ The T lymphocytes that accumulate in the lungs of patients with HP do so in an oligoclonal manner, suggesting that immune recognition of the causative antigen is responsible for the lymphocytic alveolitis.^4^

T cell-mediated immune recognition depends upon the presence of a T cell expressing a T cell receptor (TCR) with the appropriate structure to bind the complex of an antigenic peptide and the major histocompatibility complex (MHC; also known as human leukocyte antigen or HLA in humans) molecule upon which it is presented to the T cell.

The TCRs expressed by an individual’s naïve T cells are greatly diverse to allow for recognition of the diverse antigens that an individual may encounter, with the diversity resulting from the process of V(D)J recombination.^5^ V(D)J recombination is the process by which the genes encoding TCRs are assembled in each naïve T cell from the stochastic selection of variable (V), diversity (D) and joining (J) TCR gene segments in the germline.^6^ The diversity afforded by the process means that each individual’s T cell repertoire can contain in the region of 20 million unique TCR beta chain (TCRβ) amino acid sequences.^7^

Despite the richness of TCR repertoires within individuals, it is sometimes the case that the T cells bearing identical TCR amino acid sequences arise independently in multiple individuals. When this TCR sequence sharing is noted in the context of an immune response it is known as a public T cell response. Public T cell responses occur more frequently than would be explained by chance in part because V(D)J recombination is stochastic but not completely random.^8,9^ Antigen-specific public T cell responses have been observed in multiple individuals in infectious diseases, in malignancies and in autoimmune diseases.^9^

It is not known whether individuals with HP with the same causative antigen exhibit antigen-related public T cell responses. There is circumstantial evidence that this may be the case, including the observation of a shared bias in TCR gene segment expression in multiple patients with HP^10^, and the observation of beryllium-specific public T cell responses in chronic beryllium disease (an exposure-related interstitial lung disease clinically similar to HP).^11^ If this phenomenon were replicated in HP, patients with an unknown causative antigen could have the antigen identified if they shared clonally expanded TCR sequences with a patient whose antigen was known, providing that the expanded TCRs were not identifiably associated with exposures to antigens unrelated to the disease.

By reanalysing RNA sequencing data from online repositories, we found evidence of lung T cell receptor sequences shared between patients with HP. After eliminating sequences previously documented to be associated with public T cell responses to common pathogens, we identified TCR sequences plausibly specific to HP antigens, as well as candidates for the HLA alleles necessary for immune recognition of the cognate antigens. These findings suggest that an approach of surveying T cell repertoires to identify causative antigens in HP is feasible.

## Methods

### RNA sequencing data

RNA sequencing data were downloaded from the NCBI Gene Expression Omnibus (GEO; https://www.ncbi.nlm.nih.gov/geo/). The bulk of the data analysed were obtained from GEO accession GSE271789. To minimise the influence of technical factors on the composition of the reconstructed TCR repertoires, samples that were pooled prior to sequencing were excluded. Peripheral blood mononuclear cell (PBMC) samples from 12 patients with HP (all with fibrotic HP), 15 patients with idiopathic pulmonary fibrosis (IPF) and 15 healthy controls were analysed. BAL samples from 10 patients with HP (6 with fibrotic HP and 4 with non-fibrotic HP) and 10 patients with IPF were also analysed from this dataset. PBMC and BAL samples were not paired samples from the same individuals in this dataset.

Because GSE271789 did not contain BAL samples from healthy controls, data from BAL samples from 6 healthy controls were downloaded from GSE136587.

### T cell receptor repertoire reconstruction and analysis

Immune receptor repertoires were reconstructed using TRUST4 version 1.1.2, with downloaded fastq files as input.^12^ TCRβ sequences were used for analysis, with the amino acid sequences of the third complementarity determining region (CDR3) taken as the sequences in question. Incompletely sequenced CDR3β polypeptide chains were excluded, as were sequences that did not begin with an N-terminal cysteine (C) residue or end with a C-terminal phenylalanine (F) residue.

Repertoire metrics (number of clones, number of unique clonotypes) were calculated using the immunarch version 0.9.1 package in R (version 4.2.2; R Foundation for Statistical Computing). Groups were compared using Wilcoxon rank sum tests, with Bonferroni-Holm adjusted *p* < 0.05 set as the threshold for statistical significance.

Public and semi-public TCRβ sequences were identified using tcrdist3 version 0.2.2, with concatenated TCRβ repertoires as input.^13,14^ Public sequences were defined as identical TCRβ sequences occurring in more than one individual. A sequence was deemed semi-public if a sufficiently similar sequence occurred in at least one other individual. The threshold for similarity was 18 TCRdist units. TCRdist units are a weighted measure of distance between TCR sequences, with the greatest weight placed on differences in CDR3 regions and lesser weight on differences between CDR1, CDR2 and CDR2.5 sequences.^14^ Penalty scores of between 0 and 4 units for each amino acid substitution are applied (according to a BLOSUM62 substitution matrix).^15^ Where sequences are different lengths consecutive gaps are inserted into the shorter sequence at positions that minimise substitution penalties, but each gap carries the maximum penalty of four units. As an example, with CDR3 penalties carrying threefold weighting compared to other CDRs (by our criteria to determine semi-publicity), a single amino acid substitution in the CDR3 would result in two otherwise identical TCR sequences being up to 12 TCRdist units apart.

### Identification of clones under antigenic selection

TCRs were clustered into similarity neighbourhood groups, with the rationale that epitope-specific T cell responses involve expansion of both identical and highly similar T cell clones.^13^ In individual patient repertoires TCRs were placed in the same cluster if they were within 48 TCRdist units of one another, with CDR3 dissimilarities attracting a sixfold weighting in TCRdist score compared to other CDRs. This weighting allowed for neighbour dissimilarities of up to two CDR3 amino acid substitutions or gaps with identical TCR V-gene segments, or identical CDR3 sequences with differing TCR V-gene segments.^16^

The number of within-repertoire neighbours was compared to the expected number of neighbours based on a reference repertoire. 9.6 × 10^5^ TCR sequences were sampled uniformly from 8 human umbilical cord blood samples to provide an ostensibly antigen-unstimulated control reference.^16^ The cord blood samples are derived from experiments performed by Britanova and colleagues^17^, and are available through the NCBI Sequence Read Archive under accession PRJNA316572. The model comparing numbers of within-repertoire neighbours to numbers of neighbours in cord blood reference repertoires was based on the expectation of the number of neighbours following a Poisson distribution.^16,18^ Numbers of within-repertoire neighbours were considered different to expected numbers of neighbours where the *p*-values computed by the model (adjusted for multiple comparisons)^19^ were less than 0.001. Probabilities of CDR3 generation were estimated using the OLGA algorithm.^20^ Sequences occurring in expanded clones exclusive to HP patients with greater than expected numbers of within-repertoire neighbours were considered to be under antigenic selection.

### Exclusion of previously annotated TCRβ sequences

TCRβ sequences within 48 TCRdist units of entries in VDJdb^21^ (sixfold CDR3 weighting) were deemed to be related to the epitopes in the VDJdb annotations (mostly common viral and bacterial pathogens) and thus unrelated to HP. A snapshot of the VDJdb database was downloaded in May 2024 and used for the analyses.

### Prediction of HLA genotypes

HLA genotypes were predicted with T1K version 1.0.6, with fastq files as input.^22^

Predictions were made for HLA-A, HLA-B, HLA-C, HLA-DRB1, HLA-DPB1 and HLA-DQB1 genes.

## Code availability

The code to produce the analyses is available at https://github.com/wezisendama/HP_TCRsharing.

## Results

### BAL samples in HP contain greater numbers of T cell clones and unique clonotypes than control or IPF samples

As suggested by the lymphocytic alveolitis seen in HP, the TCR repertoires reconstructed from BAL samples from patients with HP contained greater numbers of clones compared to BAL samples from patients with IPF or healthy controls (**Figure 1**). Repertoires from patients with HP also contained greater numbers of unique clonotypes, suggesting concurrent expansions of multiple T cell clones in HP. There were no differences in numbers of individual clones or unique clonotypes in the repertoires reconstructed from PBMC samples, suggesting at least a partially compartmentalised pulmonary T cell immune response in HP. We therefore analysed BAL samples further.

**Figure 1.**
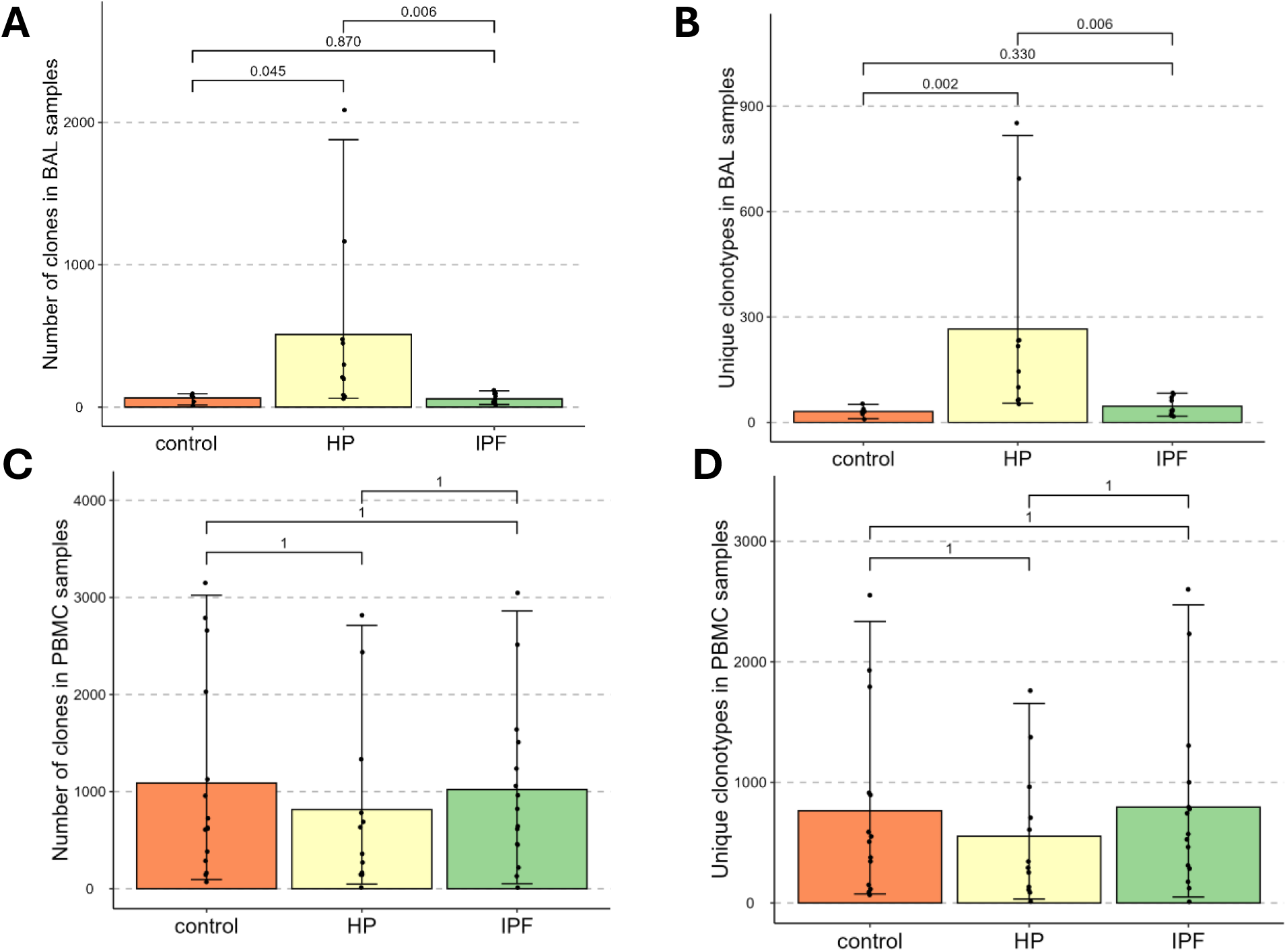
Mean number of clones and unique clonotypes in reconstructed TCRβ repertoires from BAL samples (A and B) and PBMC samples (C and D). Error bars indicate 2.5% and 97.5% quantiles. Bonferroni-Holm adjusted p values shown.

### Selected public TCRβ sequences shared between patients with HP suggest shared causative antigens

Analysis with tcrdist3 identified 15 identical CDR3β sequences shared between at least two patients with HP in BAL samples (**Table 1**). All patients with shared sequences also shared at least one HLA allele to second field resolution (**Table 2**).

**Table 1.**
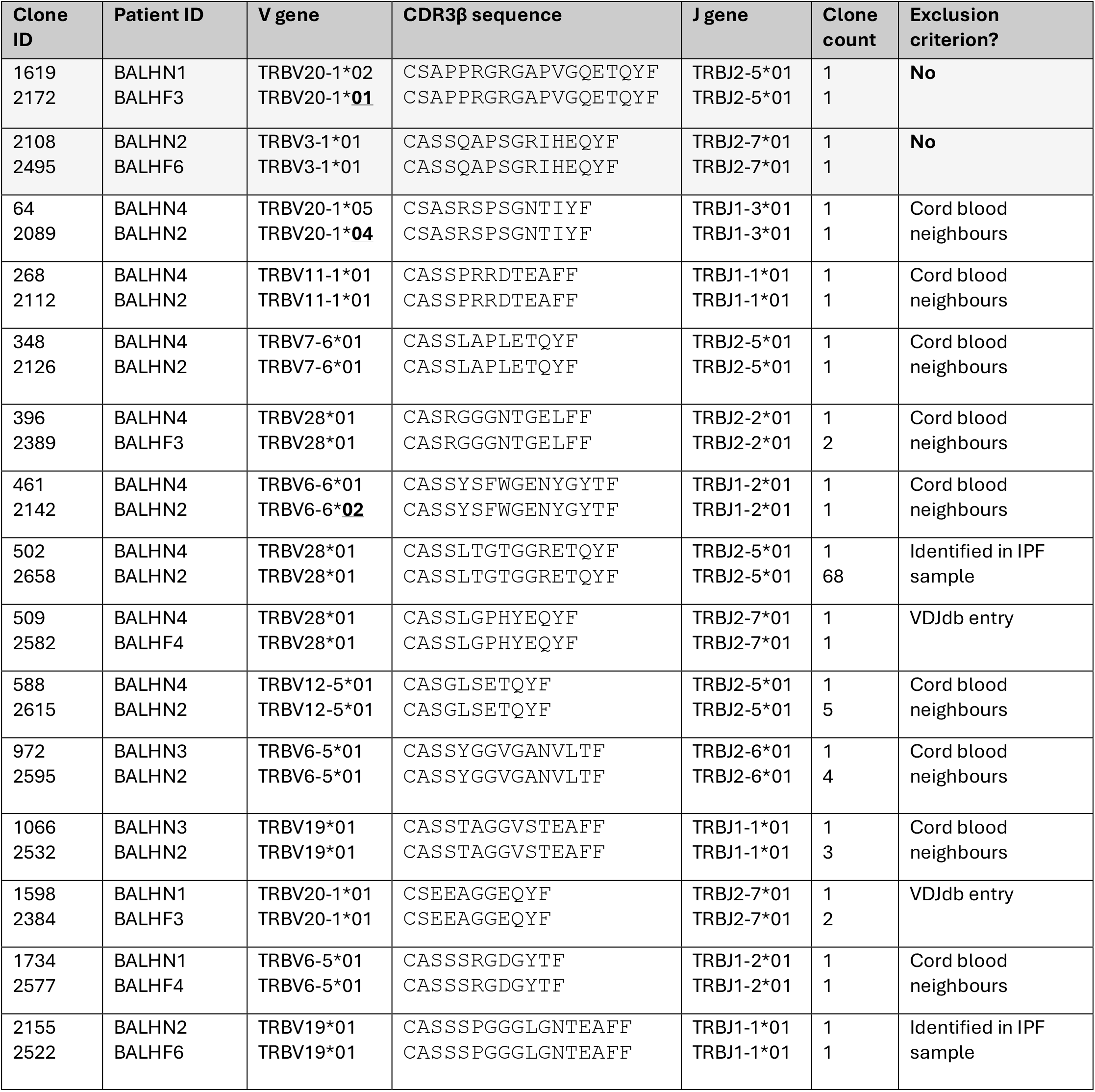
Public CDR3β sequences shared between pairs of patients with HP. Bold underlined text denotes differences in V gene segment selection despite identical CDR3β sequences. Entries in final column indicate reasons (if any) sequences could not be considered HP-related.

**Table 2.** HLA alleles of patients with HP with TCRβ repertoire overlap. HLA genotype was inferred to two-field resolution using T1K.

| Patient ID | HLA-A | HLA-B | HLA-C | HLA-DRB1 | HLA-DQB1 | HLA-DPB1 |
| --- | --- | --- | --- | --- | --- | --- |
| BALHF3 | *26:207<br>*23:01 | *07:458<br>*07:02 | *15:02<br>*15:06 | *08:77<br>*08:01 | *03:01<br>*03:72 | *1317:01Q |
| BALHF4 | *23:01<br>*31:153 | *39:150Q<br>*14:01:10 | *07:809<br>*07:906 | *04:01<br>*12:111 | *03:01<br>*03:02 | *1317:01Q |
| BALHF6 | *11:01<br>*03:312 | *38:115 | Could not be<br>inferred | *03:01<br>*04:01 | *03:01<br>*03:02 | *02:01 |
| BALHN1 | *24:02<br>*24:95 | *38:115<br>*35:471 | *04:01<br>*07:628 | *03:01<br>*08:77 | *03:01 | *02:01 |
| BALHN2 | *02:01<br>*03:350 | *13:179<br>*44:02 | *01:174<br>*02:190 | *08:01<br>*15:198 | *03:01<br>*03:72 | *02:01 |
| BALHN3 | *24:02 | *38:115 | *04:01<br>*03:02 | *04:01<br>*04:03 | *03:02 | *02:01 |
| BALHN4 | *68:01<br>*03:01 | *14:02<br>*38:115 | *05:01<br>*07:01 | *04:01<br>*03:01 | *03:01<br>*03:02 | *02:01 |

After excluding public sequences that were similar to VDJdb entries, sequences that occurred in BAL or PBMC samples from study participants who did not have HP, and sequences that had a number of within-repertoire neighbours not significantly different to the expected number of neighbours, two sequences shared between two pairs of patients with HP remained (**Table 1**, shaded rows). Although only single clones of these sequences were detected in the samples, the likelihood of identical sequences with low probabilities of generation (as determined by the OLGA estimates of generation probability as well as the absence of neighbours in cord blood samples) being detected experimentally in two individuals without clonal proliferation in the sampled repertoires is low.^23^ These sequences are therefore plausibly related to HP antigens, with the implication being that the pairs of patients share causative antigens.

13 semi-public sequences (non-identical, but within 18 TCRdist units, as described in the methods section) were identified. None of these groups of sequences met the criteria above to be considered candidate HP-related sequences.

### Expanded T cell clones with greater than expected within-group neighbours represent clones under antigenic selection

Dash and colleagues observed that TCR sub-repertoires involved in an antigen-specific immune response are composed of clusters of highly similar receptors alongside outlying receptors that are more distinct.^13^ We therefore considered whether we could use this principle to identify which clones within a patient’s repertoire could be related to the causative HP antigen even without evidence of sequence sharing with another patient with HP. Bearing in mind that sequences with a higher probability of being generated by V(D)J recombination are more likely to cluster even without the influence of antigen recognition^24^, we also compared each sequence’s number of within-repertoire neighbours to its expected number of neighbours (using cord blood samples as the antigen-naïve reference).

After excluding sequences within 48 TCRdist units of VDJdb entries and sequences that also occurred in samples from participants who did not have HP, several examples were identified of clones in the BAL repertoires of individual patients that may be related to causative antigens in HP (**Table 3**).

**Table 3.**
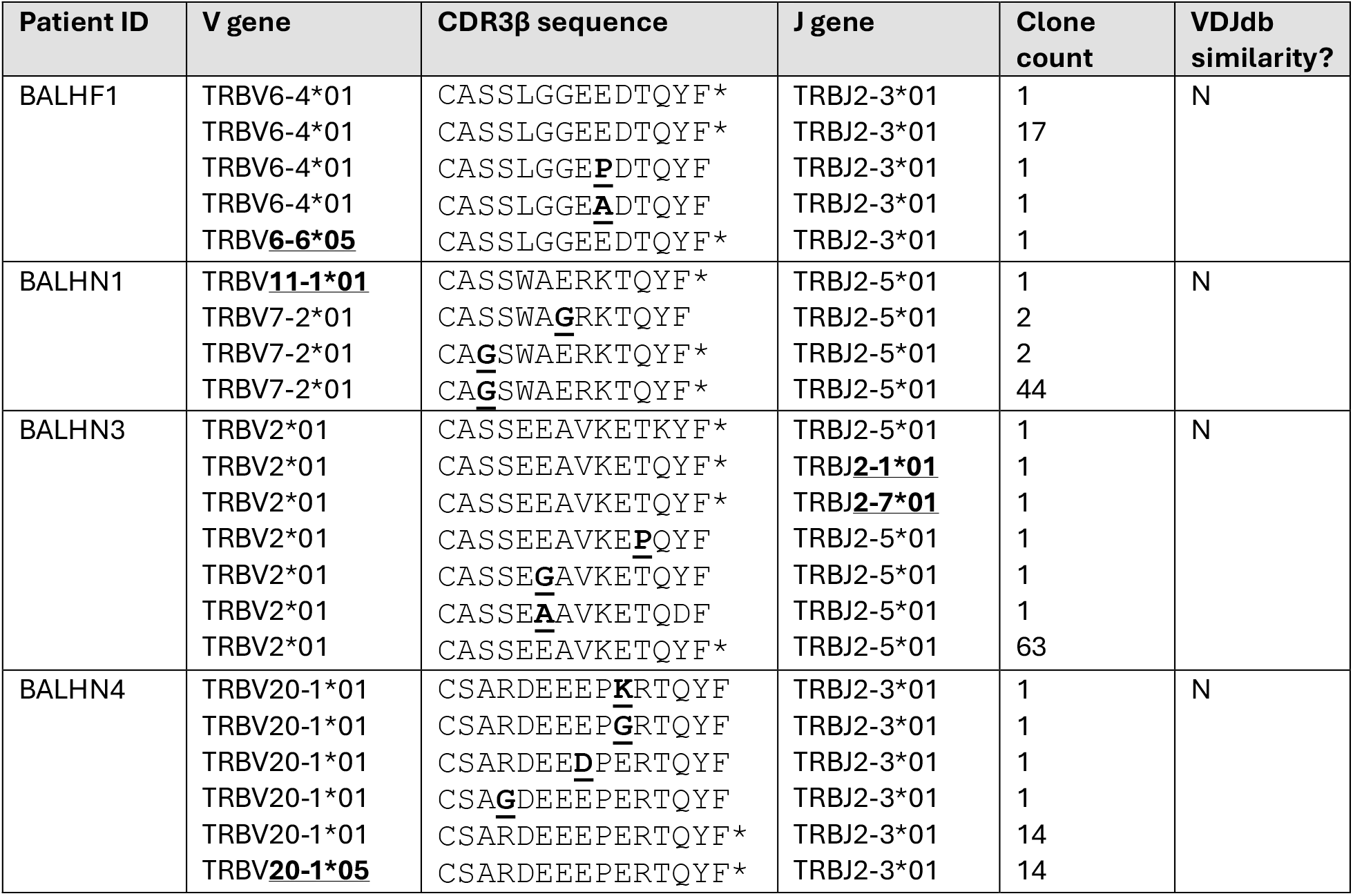
Clusters of highly similar T cell clones presumed to be under antigenic selection. Bold underlined text indicates differing V gene selection and CDR3β amino acid substitution. Asterisks denote identical CDR3β amino acid sequence with differing CDR3β nucleotide sequence or V gene.

## Discussion

We present evidence of TCR sharing in HP that is likely to be related to the causative HP antigens in the patients sharing the receptor sequences. To our knowledge, TCR sharing in HP has not been reported previously. Antigen identification in HP remains a challenge, but we are hopeful that the principle established here might allow identification of a patient’s causative antigen where HP-related TCRs are shared with a patient whose antigen is known.

We also identify clusters of TCR sequences in individual patients that are likely to be related to disease in HP. While we cannot present evidence of these sequences being shared with other patients, we suggest that there is value in establishing a database similar to VDJdb that annotates such sequences with the patients’ causative antigens and HLA genotype where known. Where the sequences are subsequently identified in other patients with HP sharing the same antigen (and at least one HLA allele) an annotation denoting the level of confidence in the association can be updated.

Conversely, where the sequence is confirmed experimentally to be associated with a different pathogen the database entry may be removed. Antigen identification could therefore be aided by cross-references against the database. Open access of the database could allow contributions from clinicians and researchers globally, maximising its utility.

There are limitations in our study. The absence of BAL samples from control participants in the GSE271789 dataset means that data from separate experiments had to be considered. It appeared that the T cell repertoires reconstructed from the healthy control BAL samples from GSE136587 were of similar size to the repertoires from patients with IPF from GSE271789. This is expected and gives some confidence in the validity of the control data given that there are similar numbers of lymphocytes per unit of BAL volume in patients with IPF and healthy controls^25^, but we cannot exclude the impact of technical differences between the experiments that yielded the repository data.

Another limitation was the absence of paired BAL and PBMC samples from participants in the GSE271789 dataset. This could have increased the confidence in our finding of a compartmentalised immune response in HP by allowing clonotypes to be tracked between tissue compartments. There is prior evidence that supports a proportionally greater T cell response in the lung compared to peripheral blood in HP including the observation of a lung-specific lymphocyte expansion that abates when the causative antigen is removed^4^, and our own evidence of lung-resident T cells providing an appreciable portion of the HP immune response.^26^ These study results support our findings from the comparisons of BAL and PBMC repertoire sizes despite us not having access to paired samples.

While VDJdb represents a vast database of experimentally confirmed TCR specificities, by definition it cannot be complete. We aimed to account for this by excluding TCR sequences from consideration as HP-related if they were within a liberally defined distance of VDJdb entries, and then by considering shared TCRs that have a low probability of generation and did not occur in non-HP samples. While we therefore cannot completely exclude the possibility that the pairs of individuals share the unexpected TCRs because of responses to an HP-unrelated epitope, we believe that a lung compartment TCR without neighbours in an antigen-naïve reference sample shared by two individuals with the same lung disease is unlikely to be explained by chance.

Our work here establishes the feasibility of using T cell repertoires to identify antigens in HP, and our use of publicly available RNA sequencing data is a strength as it implies that it can be done at relatively low-cost using samples collected for other indications. A low-cost method to identify antigens in HP has the potential to improve outcomes in the disease, and future studies should aim to establish the clinical utility of such an approach in the management of the disease.

